# Selection at behavioral, developmental and metabolic genes is associated with the northward expansion of a successful tropical colonizer

**DOI:** 10.1101/478214

**Authors:** Yann Bourgeois, Stéphane Boissinot

**Affiliations:** New York University Abu Dhabi, Saadiyat Island Campus, Abu Dhabi, United Arab Emirates

## Abstract

What makes a species able to colonize novel environments? This question is key to understand the dynamics of adaptive radiations and ecological niche shifts, but the mechanisms that underlie expansion into novel habitats remain poorly understood at a genomic scale. Lizards from the genus *Anolis* are typically tropical and the green anole *(Anolis carolinensis)* constitutes an exception since it expanded into temperate North America from subtropical Florida. Thus, we used the green anole as a model to investigate signatures of selection associated with colonization of a new environment, namely temperate North America. To this end, we analyzed 29 whole genome sequences, representing the entire genetic diversity of the species. We used a combination of recent methods to quantify both positive and balancing selection in northern populations, including F_ST_ outlier methods, machine learning and ancestral recombination graphs. We naively scanned for genes of interest and assessed the overlap between multiple tests. Strikingly, we identified many genes involved in behavior, suggesting that the recent successful colonization of northern environments may have been linked to behavioral shifts as well as physiological adaptation. These results were robust to recombination, gene length and clustering. Using a candidate genes strategy, we determined that genes involved in response to cold or behavior displayed more frequently signals of selection, while controlling for local recombination rate and gene length. In addition, we found signatures of balancing selection at immune genes in all investigated genetic groups, but also at genes involved in neuronal and anatomical development in Florida.

## Introduction

Colonization and adaptation to new environments can result in remarkable phenotypic divergence and is of great significance at a time of acute climate change. It is therefore critical to assess the mechanisms of adaptation to predict the resilience of species to change or their ability to colonize and invade novel habitats. However, the genetic bases of adaptation and their evolutionary causes remain unclear with few exceptions (Abzhanov et al. 2008; Steiner et al. 2009; Hohenlohe et al. 2010; Weber et al. 2013; Tiffin and Ross-Ibarra 2014). This raises the question of the nature of the proximate mechanisms by which populations and species overcome their tendency to remain in similar environments. Such a question can only be addressed in a broad phylogenetic context since organisms differ in their intrinsic ability to colonize new habitats. Understanding the evolution of phenotypic innovations has led to extensive studies on the behavioral and physiological changes associated with adaptation to new environments (Butler et al. 2002; Pincheira-Donoso et al. 2008; Campbell-Staton et al. 2017; Lapiedra et al. 2018). Ultimately, a comprehensive framework would address both environmental and genetic causes of adaptation (Mayr 1961). Advances in high throughput sequencing now provide a way to use whole genomes to investigate the interactions between molecular diversity and evolutionary processes.

From a genomic perspective, adaptation to new biotic and abiotic constraints should lead to signatures of positive selection that are scattered across the genome (Qvarnström and Bailey 2009; Seehausen et al. 2014). Identifying loci that are important in adaptation and divergence remains a major technical challenge of population genomics, which has led to the extensive development of method targeting signatures of natural selection across genomes. Most of these genome-wide scans for selection (GWSS) were initially dedicated at identifying genomic signatures of positive selection in humans (Haasl and Payseur 2016). These approaches have subsequently led to dramatic improvements in our understanding of adaptation in other model species such as *Drosophila* (Garud et al. 2015; Sheehan and Song 2016), *Arabidopsis* (Kubota et al. 2015; Alonso-Blanco et al. 2016), or maize (Beissinger et al. 2014; Tiffin and Ross-Ibarra 2014). More recent studies have used these methods in non-model species, contributing to a better understanding of the links between genetic variation, adaptation and speciation (Haasl and Payseur 2016; McKinney et al. 2017). Genome scans are no longer limited to recent positive selection but can also track signatures of balancing selection in genomes, a type of selection that is commonly found at immune and resistance genes (Sommer 2005; Tellier and Brown 2011; Bento et al. 2017). Although this aspect may have been overlooked (Haasl and Payseur 2016; Siewert and Voight 2017), finding signatures of balancing selection holds the potential to elucidate the mechanisms that explain the long-term persistence of alleles in populations.

Anoles are a neotropical group of squamates that diversified during the Cenozoic, and constitute a model system for understanding adaptation and speciation in ectotherms (Losos et al. 2004; Glor et al. 2005; Losos 2009; Kolbe et al. 2017; Lapiedra et al. 2018). Anoles display an extraordinary physiological, morphological and behavioral diversity (Lailvaux et al. 2004; Glor et al. 2005; Campbell-Staton et al. 2018; Lapiedra et al. 2018; Losos 2009). This diversity is reflected in genomic variation across anole species, with clear signatures of positive selection at developmental genes (Tollis et al. 2018). While this suggests recurrent selection at these loci over macroevolutionary scales, a thorough investigation at the whole genome level has never been led on more recent adaptation (but see (Campbell-Staton et al. 2016; Prates et al. 2018) for studies using reduced representation methods). In this context, the green anole *(Anolis carolinensis)* is an ideal system to test how adaptation to new environment impacts genome variation since it is the only species of the genus to have expanded into temperate territories (Campbell-Staton et al. 2012, 2017, 2018; Goodman et al. 2013; Manthey et al. 2016; Ruggiero et al. 2017; Tollis et al. 2018). Indeed, ectotherms face unique challenges to adapt to cold climates, and increase their fitness through behavioral, physiological and morphological means (Blouin-Demers et al. 2003; Bouazza et al. 2016; Campbell-Staton et al. 2018).

The ancestor of the green anole originally colonized Florida from Cuba between 6 and 12 million years ago (Glor et al. 2005). A first step of divergence occurred in Florida between 3 and 2 mya (Figure 1) and was presumably caused by sea level changes (Tollis and Boissinot 2014), which resulted in the formation of archipelagos in place of what is now Florida. This process produced three distinct genetic clusters in Florida, the North-Eastern Florida clade (NEF), the North¬Western Florida clade (NWF) and the South Florida clade (SF). The ancestral population of lizards now living in temperate territories diverged from the NEF clade approximately 1 mya. This divergence was followed by expansion northwards from Florida along the Atlantic seaboard and west across the Gulf Coastal Plain over the last 100,000 years (Manthey et al. 2016;

**Figure 1.**
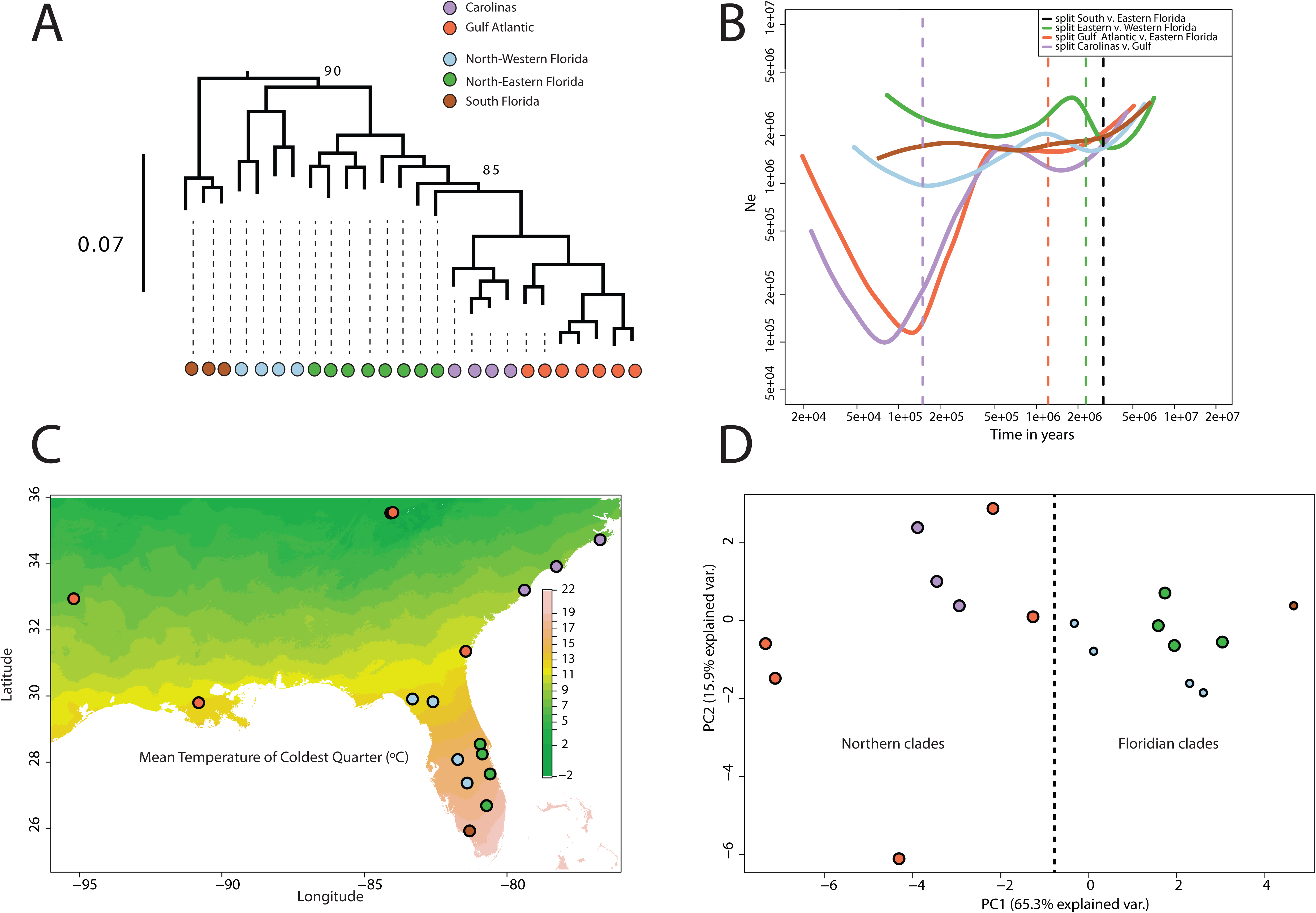
Summary of population structure and environmental variation in green anoles. A: RAxML phylogeny on one million random SNPs (Bourgeois et al. 2018a). B: Demographic evolution of the five genetic clusters of green anoles reconstructed by SMC++. C: Sampling locations used in this study. Units for temperature are in tenth of Celsius degrees. D: PCA over environmental variables (BIOCLIM data) for the locations used in this study. Larger dots highlight the northern clades (GA and CA) and their sister Floridian clade (NEF).

Bourgeois et al. 2018a). This led to the emergence of the two current northern clades, Gulf-Atlantic (GA) and Carolinas (CA). Lizards from these northern populations are bigger than their Floridian counterparts (Goodman et al. 2013), have likely encountered environmental conditions never experienced by any other anole species during northwards colonization (Campbell-Staton et al. 2016) and display physiological adaptation to cold (Campbell-Staton et al. 2018). A robust annotated reference genome is available (Alföldi et al. 2011), and previous population genomics studies (Campbell-Staton et al. 2012; Tollis et al. 2012; Tollis and Boissinot 2014; Manthey et al. 2016) provide the necessary background to perform rigorous tests for selection that take into account genome structure, variable recombination rates and demography.

In this study, we investigate the genetic bases of adaptation associated with the colonization of a novel environment using a set of 27 whole genomes sampled across the species range (Figure 2). We use a combination of methods (differentiation outliers, population branch statistic, machine learning, ancestral recombination graphs and allele correlations) to quantify both positive and balancing selection in northern populations while controlling for the effects of demography. We interpret these GWSS in two ways (Figure 2). First, we naively scan for genes and GO terms of interest in the set of candidates for selection, and assess the overlap between multiple tests. We further adopt a candidate genes strategy to test whether genes involved in development, physiology and behavior display more often signals of selection, while controlling for local recombination rate and gene length. Our study provides important results to understand the genetic causes of adaptation over recent evolutionary timescales.

**Figure 2.**
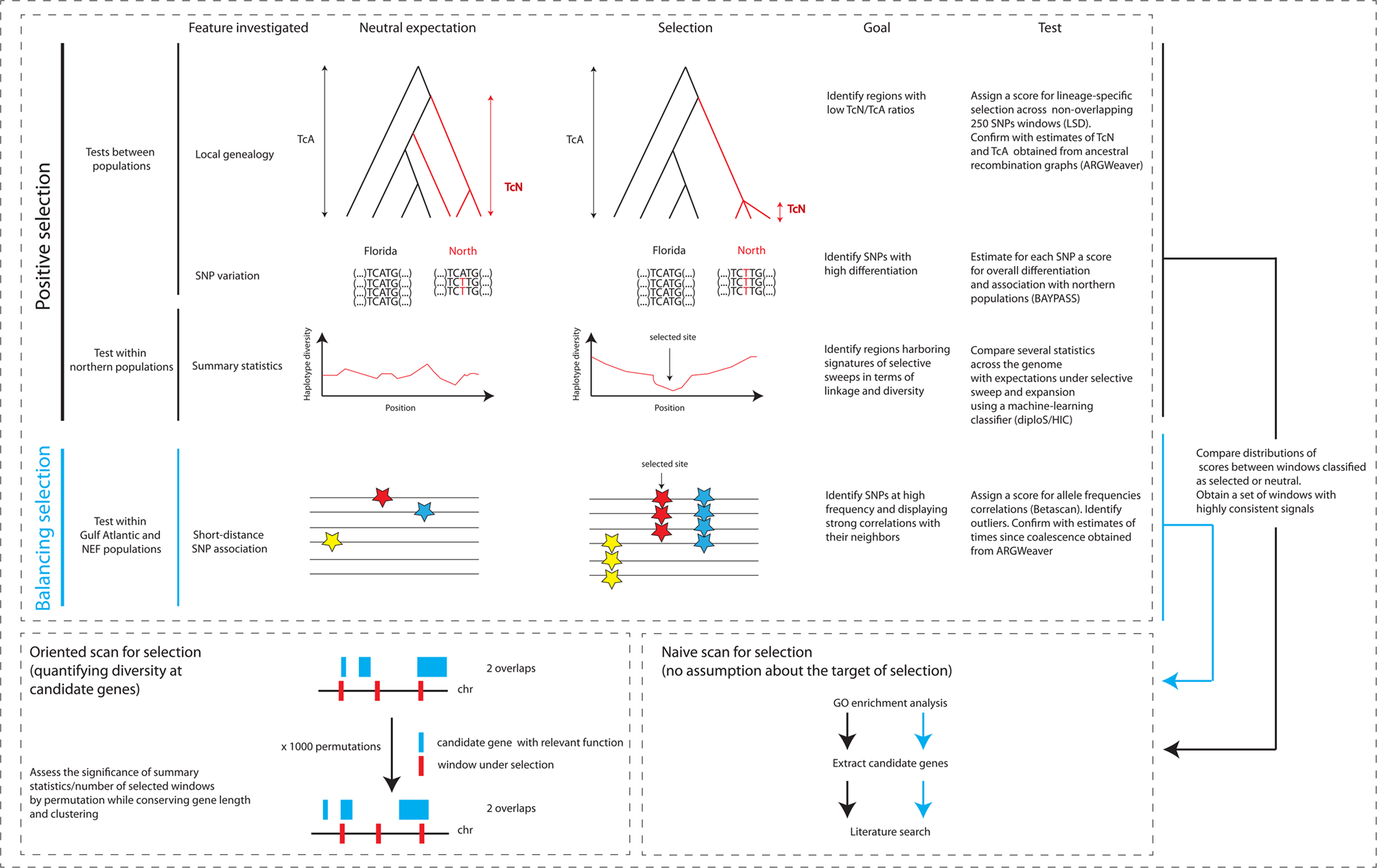
Flowchart of the methods used in this study. TcA: Time since coalescence in all individuals/populations. TcN: Time since coalescence in Northern populations.

## Results

### Naive scan for genes under positive selection

Selective sweeps result in spatial variation in the allele frequency spectrum and linkage disequilibrium around the selected locus (Figure 2). We scanned the genome of the two Northern populations for signatures of selection using diploS/HIC, a classifier trained over simulated datasets that incorporate demography and selection. The method further attempts to classify whether selection occurred on a *de novo* mutation (“hard sweep”) or on standing variation (“soft sweep”). Simulated datasets are split into subwindows that are described by a set of 12 summary statistics (Kern and Schrider 2018) recapitulating the allele frequency spectrum or linkage disequilibrium. In the case of selection, simulated windows where the selected site lies in the central subwindows are considered hard or soft sweeps examples, while other windows are considered linked-hard or linked-soft examples. This set of simulated datasets is then used to train a supervised machine-learning algorithm that uses the spatial organization of summary statistics to differentiate between each category. Predictions on the actual genomic dataset are then performed over subwindows.

Three thousand nine hundred twenty-eight 30kb windows were classified as selective sweeps with a false positive rate of 5% and a true positive rate of 55-60% (Sup. Table 1). The algorithm was not able to reliably discriminate between hard and soft sweeps. The total number of 30kb windows included in this analysis was 45,717, due to the exclusion of scaffolds < 3.3Mb. Outlier windows covered 2027 genes. To explore the function of these loci, we ran a GO term enrichment test for biological process (Sup. Table 2). The first 50 most significant terms were related to metabolism and regulation (e.g. positive regulation of catalytic activity, p= 4.61.10^−5^; regulation of metabolic process, p=3.8.10^−4^) as well as behavior and nervous system (e.g. cognition, p=1.44.10^−3^; behavior, p=3.21.10^−3^).

**Table 1.**
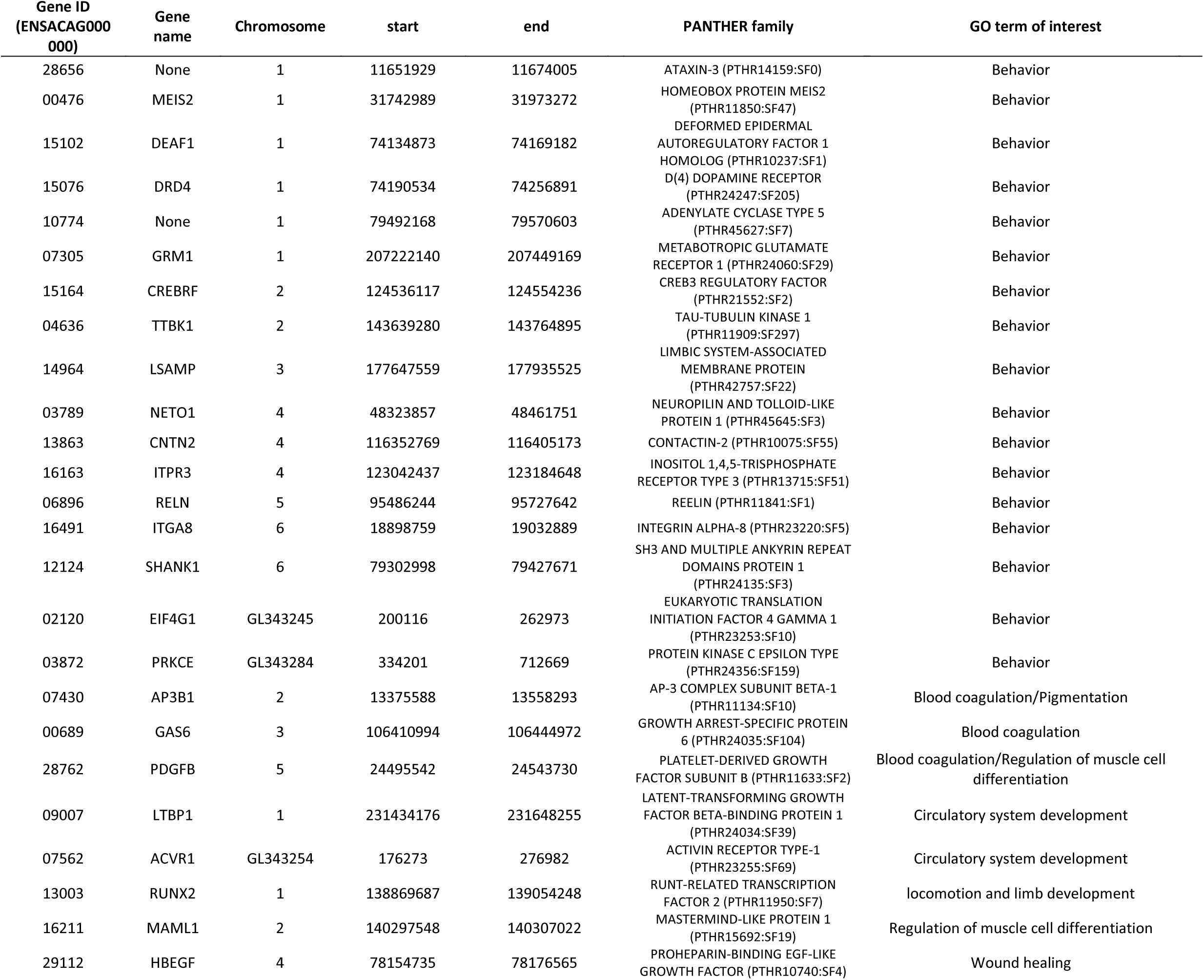
Candidate genes for positive selection. All genes overlapped with windows classified by diploS/HIC as selective sweeps that were in the top 10^th^ percentile for LSD and median eBPis scores.

**Table 2.**
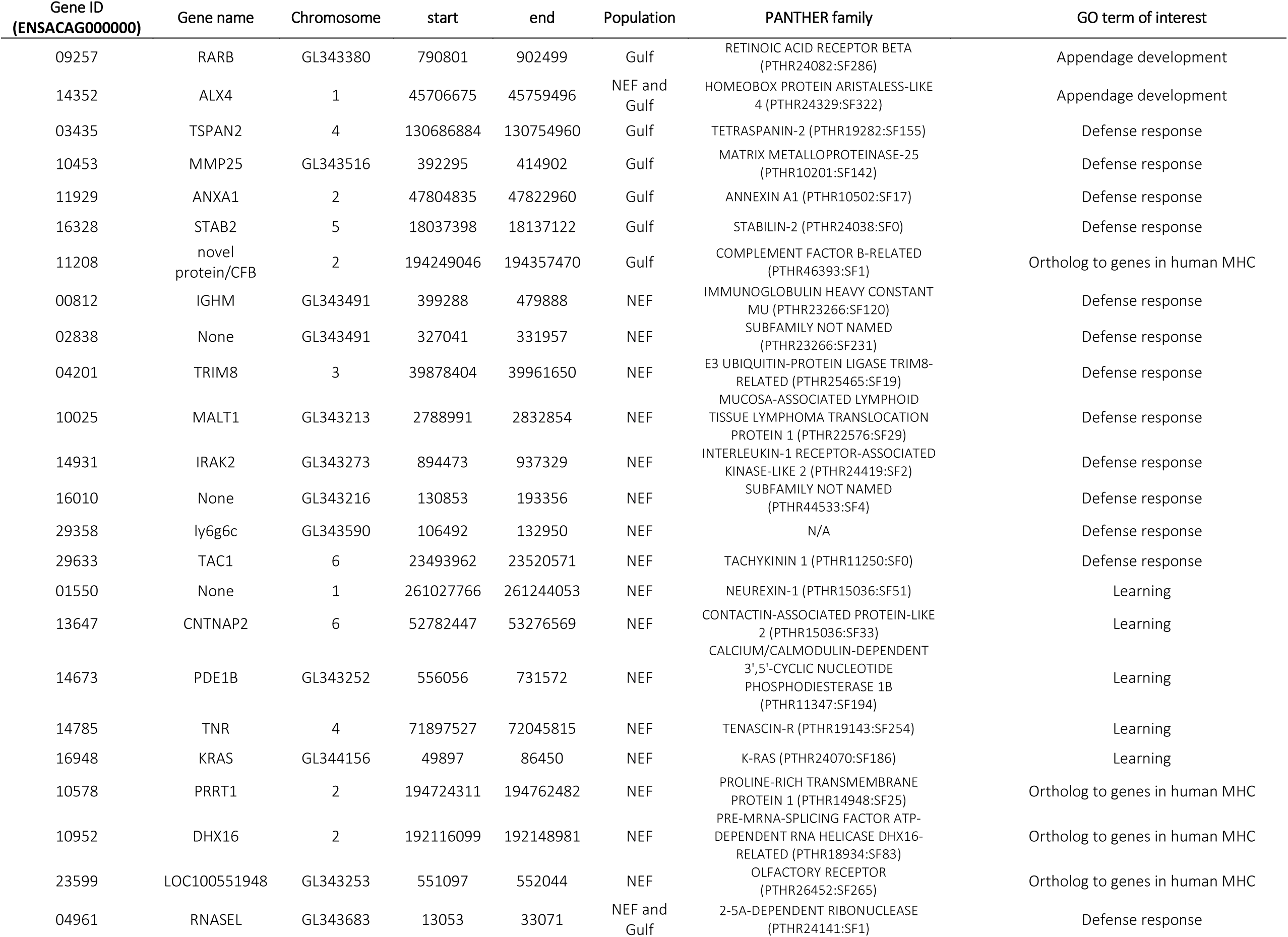
Candidate genes for long-term balancing selection in NEF and GA clusters.

To detect genomic regions displaying signatures of positive selection in the branch leading to northern clusters, we used the LSD algorithm (Librado and Orlando 2018). This method compares the levels of exclusively shared differences between populations across genomic windows. The LSD score should be maximized when the local tree displays evidence for recent coalescence within the focal populations and high differentiation compared to the rest of the genome (Figure 2). We applied the LSD test over 50,996 windows encompassing 1,250 SNPs each (~30kb per window, 1.69 Gb covered) to detect genomic regions displaying evidence for recent selective sweeps along the branch leading to Northern populations. The 1,000 top-ranking windows for LSD score had a minimal score of 0.779 and covered 834 genes. The top 50 enriched GO terms (Sup. Table 3) were related to regulation (e.g. DNA packaging, p=1.01.10^−4^), metabolism (e.g. metabolic process, p=1.23.10^−3^), gonad development (regulation of male gonad development, p=3.10^−3^) and behavior (p=8.13.10^−3^).

**Table 3.** Z-scores comparing the distance between observed average recombination rate, eBPis, LSD scores and number of overlaps with windows with consistent signals of positive selection with 1000 permuted datasets. For behavior and cold response genes, a second test was run removing genes falling in the bottom 25% recombination rate of all genes. One-tailed p-values for higher scores or lower recombination rate: *, p<0.05; **, p<0.01; ***, p<0.001.

| Gene category | GO term/Reference | Average recombination rate | Average eBPis | Average LSD | # overlaps with three methods |
| --- | --- | --- | --- | --- | --- |
| Behavior (N=295) | GO:0007610 | -1.23/0.35 | **2.788\*\*/1.776\*** | **4.334\*\*\*/2.183\*** | **5.217\*\*\*/3.213\*\*** |
| Immune response (N=450) | GO:0006955 | -1.469 | 0.889 | 1.565 | 1.522 |
| Somitogenesis and limb development (N=166) | GO:0001756 ; GO:0060173 | 0.534 | -1.649 | 0.781 | -0.971 |
| Cold response genes (N=1375) | Campbell-Staton et al. 2018 | **-4.202 \*\*\*/-0.505** | **2.579\*\*/1.963\*** | **4.57\*\*\*/2.022\*** | 1.554/0.222 |
| Reproduction (N=479) | GO:0000003 | **-2.851 \*\*** | 1.489 | 1.1 | 1.541 |

Methods scanning windows across genomes that are not fully assembled cannot use short genomic scaffolds since they are not informative enough. Given the rather high number of short scaffolds in the green anole genome, we used the approach implemented in BAYPASS (Gautier 2015) that is aimed at detecting overtly differentiated SNPs (Figure 2). Our aim was to identify SNPs with high differentiation and discriminating between Northern and Floridian populations. Seven hundred twenty-six 5kb windows harboring at least three outlier SNPs were identified over a total of 249,351. These outlier windows encompassed 284 genes. The top 50 enriched GO terms (Sup. Table 4) were related to signaling (p=1.93.10^−5^), metabolism (e.g. cyclic nucleotide metabolic process, p=4.47.10^−4^), cellular response (e.g. response to platelet aggregation inhibitor, p=1.66.10^−4^) and behavior (locomotory exploration behavior, p=5.67.10^−3^).

To evaluate the overlap and consistency between these methods, we examined the distribution of statistics for selection in regions classified as sweeps or neutral by diploS/HIC. To facilitate comparisons between LSD and eBPis, we computed the median eBPis score over the same windows as LSD. Candidate windows for selection (hard and soft sweeps combined) harbored an excess of high LSD and median eBPis scores compared to windows classified as neutral, the effect being particularly clear for the LSD statistics (Figure 3). Based on these distributions, we extracted a set of windows with consistent signals of selection. These 1,250 SNPs windows had a minimum overlap of 50% with windows classified as selected by diploS/HIC and belonged to the top 10% for median eBPis and LSD scores. GO term enrichment tests (Sup. Table 5) over the set of 402 candidate genes covered by these windows highlighted functions related to metabolism (e.g. regulation of metabolic process, p=4.44.10^−4^), behavior (p=6.32.10^−4^) and cognition (p=3.66.10^−3^).

**Figure 3.**
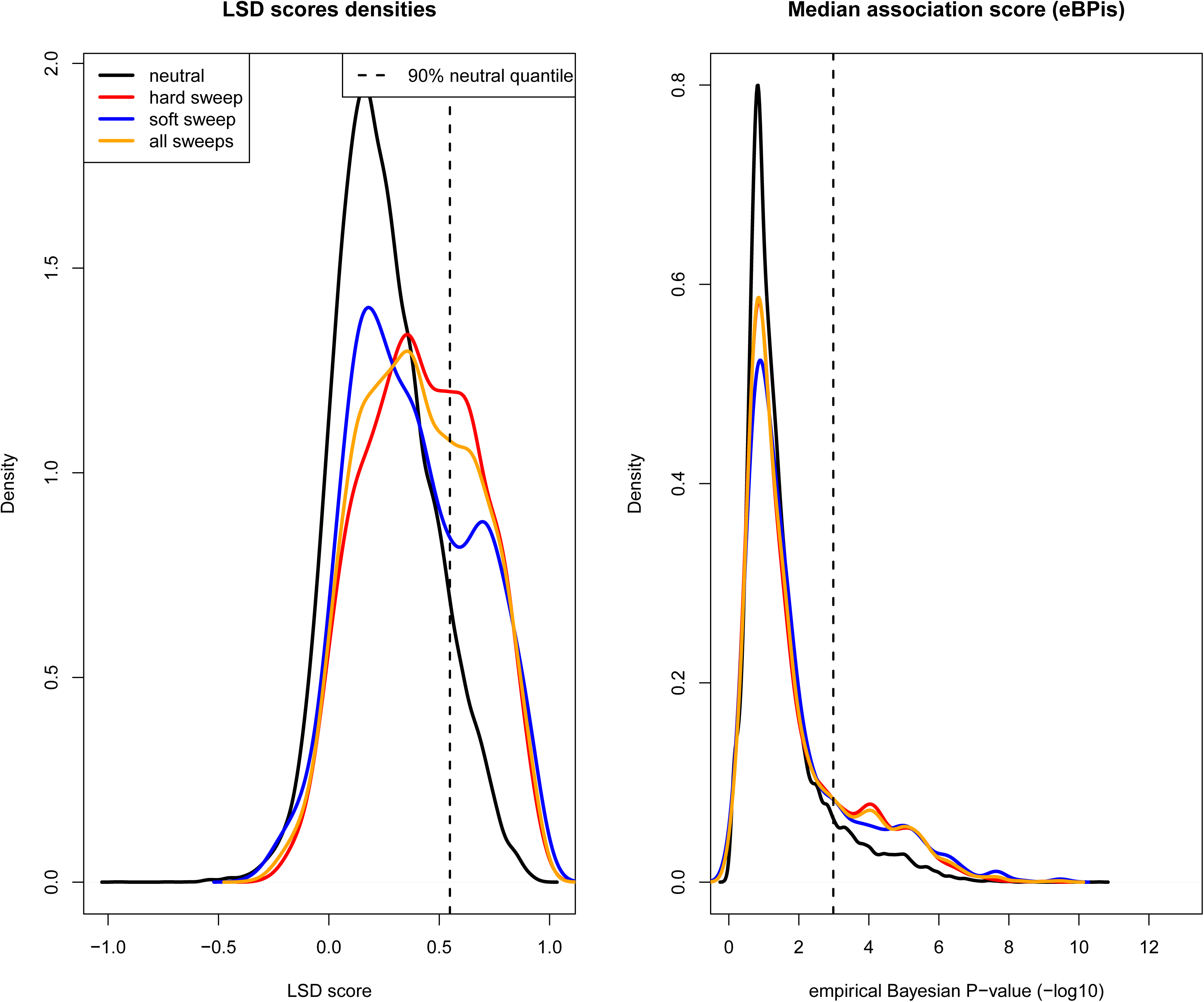
Comparison of LSD and eBPis scores in windows overlapping 30kb intervals classified as neutral or selected by diploS/HIC.

We focused on GO categories that are *a priori* relevant either because of the ecology of green anoles or more generally relevant to anoles diversification (Losos 2009; Tollis et al. 2018). We retrieved a list of genes involved in limb development, behavior, coagulation, or vascular system development (Table 1, Figure 4). We did not notice any strong clustering of candidate genes assigned to the GO category “behavior”, suggesting that their overrepresentation in the set of selected genes was not due to shared overlaps with candidate windows for selection. In general, signals of selection displayed consistent spatial patterns, with eBPis and LSD scores rising at the vicinity of windows classified as selected by diploS/HIC (Figure 4).

**Figure 4.**
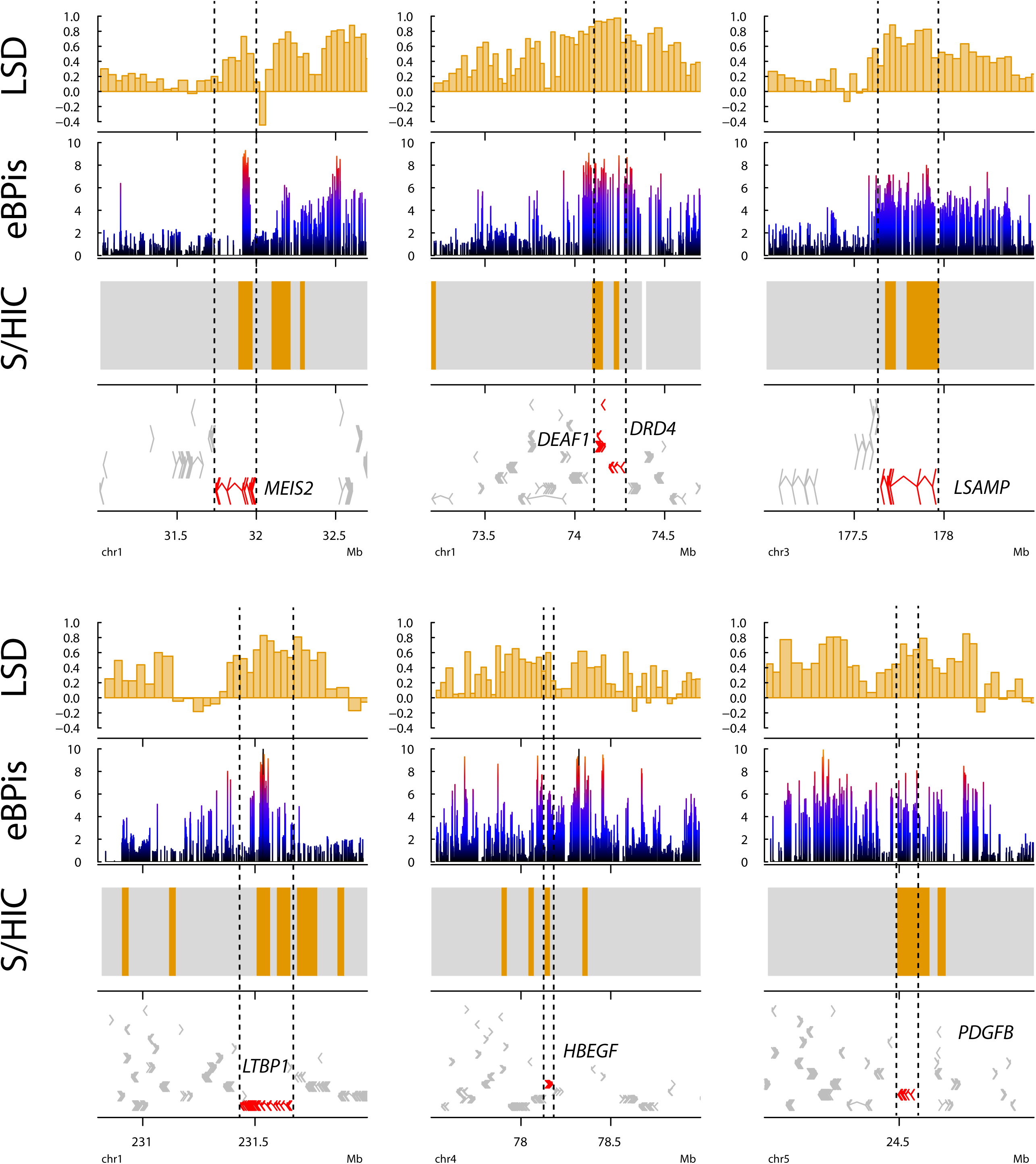
Plots of statistics for positive selection at seven candidate genes. For the diploS/HIC track, orange windows are classified by selected, grey windows are classified as neutral or linked.

### Naive scan for genes under balancing selection

Next, we scanned for genes under balancing selection in Gulf Atlantic and its source cluster, North Eastern Florida. Long-term balancing selection leads to the build-up of allelic classes and leads to strong correlation in frequencies between SNPs close from sites under long-term balancing selection (Figure 2). This pattern can be captured by a recently developed statistics, β (Siewert and Voight 2017). We computed β scores over SNPs in non-overlapping 5 kb windows with Betascan. We retrieved 244 and 577 windows (over 324,131 and 333,258) with at least three outlier SNPs in the top 0.1% β score in GA and NEF respectively. These outlier windows overlapped 89 (GA) and 191 (NEF) genes. In GA, the 50 first most significant GO terms were related to immunity (Sup. Table 6, e.g. granulocyte migration, p=2.73.10^−4^; regulation of interleukin-2 production, p=7.75.10^−3^) and developmental processes (e.g. developmental process, p=3.18.10^−3^; embryonic hindlimb morphogenesis, p=5.09.10^−3^). In NEF, GO terms related to development displayed extremely low p-values (Sup. Table 7, e.g. system development, p=3.46.10^−6^; nervous system development, p=5.11.10^−6^; anatomical structure development, p=1.43.10^−4^). GO terms related to learning and behavior also belonged to the top 50 most significant terms with 5 genes under the term “learning” (p=1.05.10^−3^). A more detailed list of relevant candidate genes is provided in Table 2 and Figure 5.

**Figure 5.**
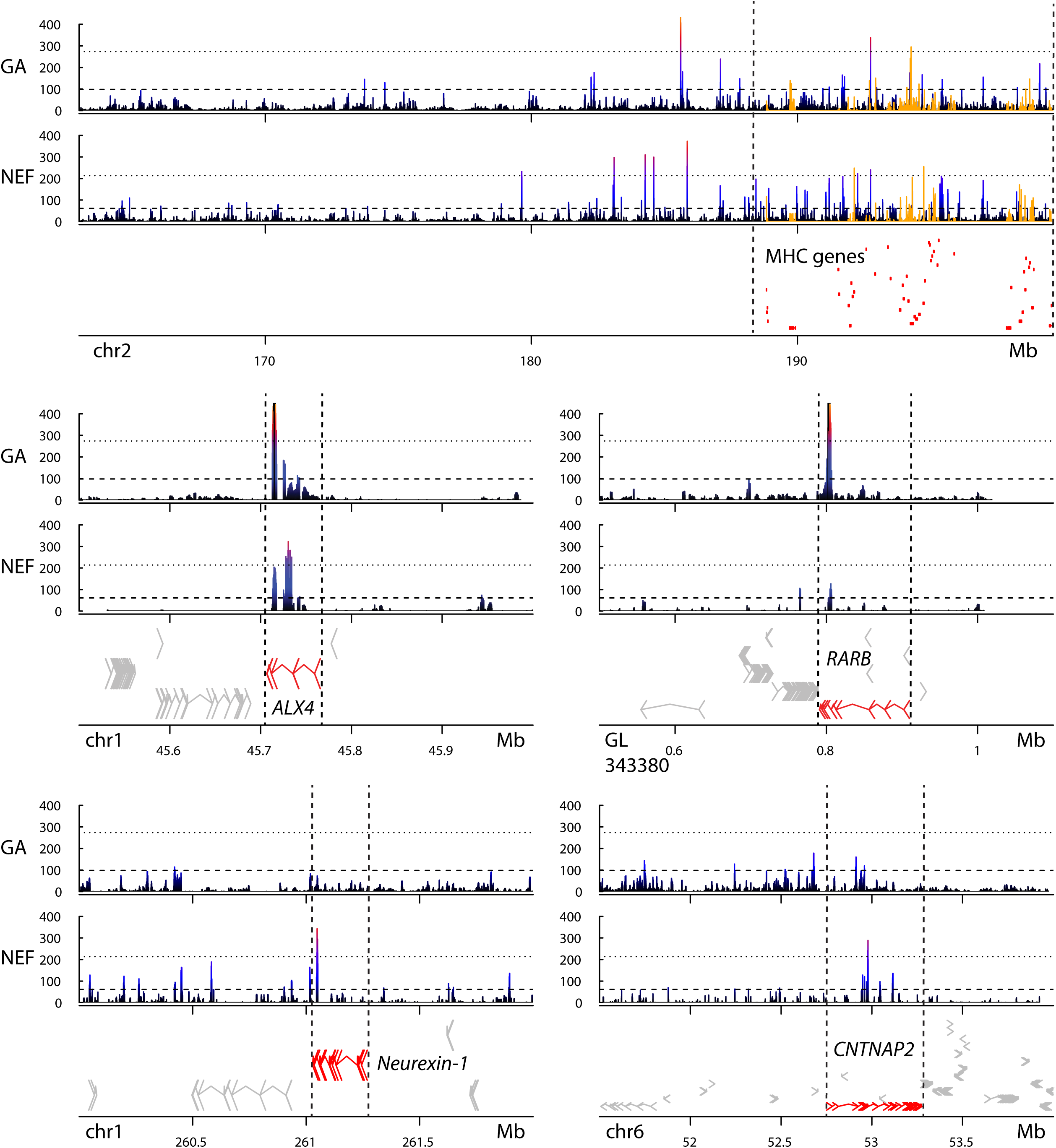
Plots of β scores in NEF and GA at the vicinity of the region homolog to human MHC (top panel) and four other candidate genes (bottom panels). For better clarity, only genes homolog to human MHC are displayed in the top panel. Orange peaks correspond to SNPs found at the vicinity (+/-5kb) of human MHC homologs.

A classic example of balancing selection includes the MHC genes in Vertebrates. We examined the distribution of β scores along a region harboring a high concentration of genes homolog to human MHC on chromosome 2 (Figure 5). This region harbored many SNPs overlapping with MHC homologs with β scores in the top 0.1 and 1% for both NEF and GA clusters.

To control for unknown paralogs inflating the number of intermediate frequency alleles and therefore β scores, we investigated whether outlier SNPs lying in candidate genes displayed any significant excess of heterozygotes in GA, NEF and across all populations. Less than 2% of these variants had p-values below 0.01 across all tests, making ancient gene duplication an unlikely explanation for the observed pattern.

### Selection and time since coalescence

The coalescence time for the set of genes with consistent signals across all three methods for detecting positive selection was strongly reduced compared to genomic background in Northern clades, but did not display any significant reduction at the whole species scale (Figure 6). This is consistent with positive selection and hitchhiking effects reducing coalescence times around loci under positive selection in Northern populations (Figure 2).

**Figure 6.**
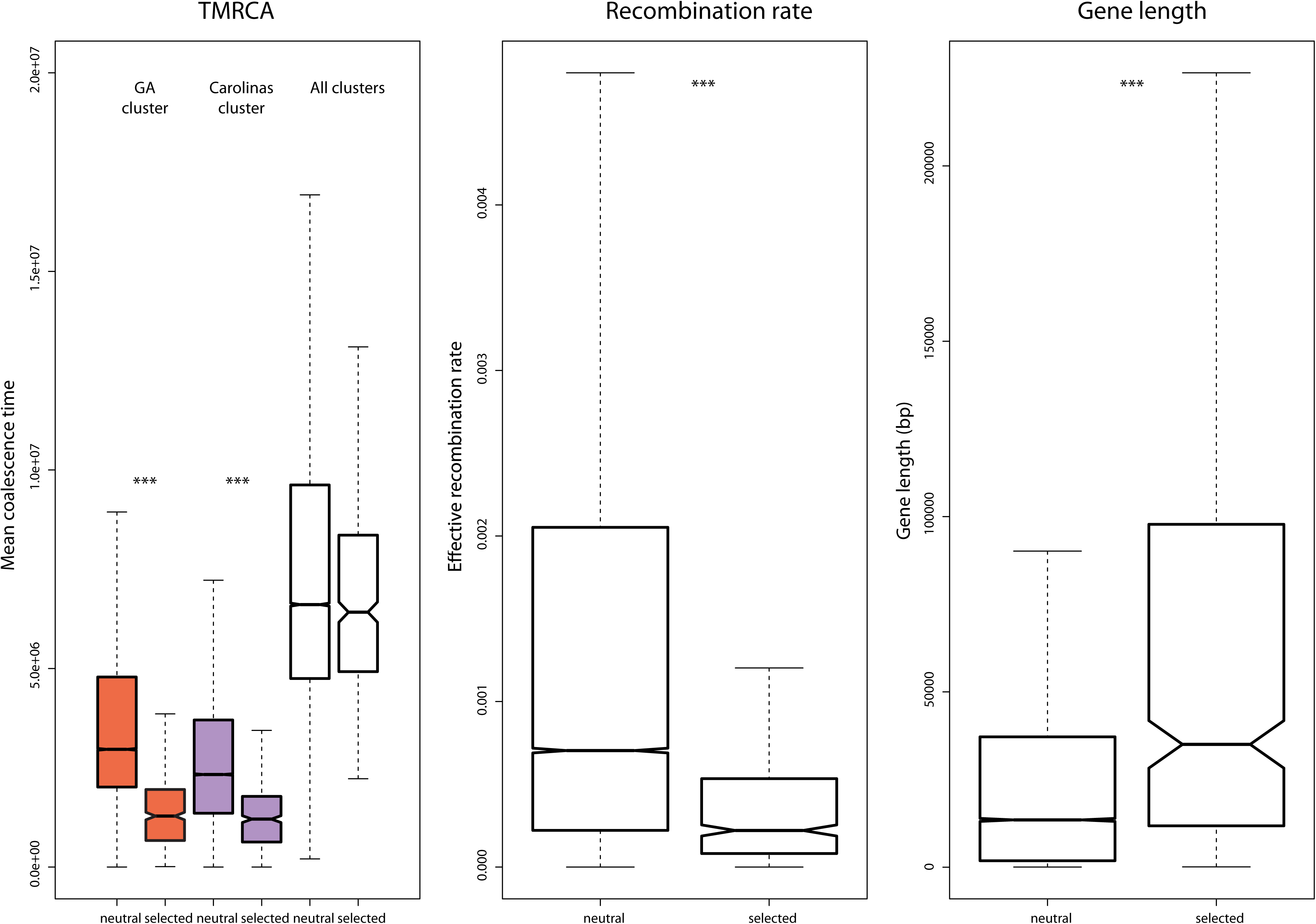
Properties of windows under positive selection in terms of coalescence time, gene length and effective recombination rate (calculated in NEF). ***: p-value<0.001, Wilcoxon tests. For gene length, only genes overlapping windows that were common between all tests were included in the background set.

Genes under long-term balancing selection should display older coalescent times (Charlesworth 2006). The results from Betascan were consistent with this expectation, with outlier windows that displayed significantly longer coalescence times compared to genomic background. These longer times are typical under balancing selection, as alleles are maintained over long periods of time (Charlesworth 2006). We assessed whether diversity at genes under balancing selection was maintained despite the expansion event in northern populations by examining coalescence times in GA for outliers identified in NEF. The loss of diversity in Gulf Atlantic compared to its sister clade was reflected in the reduced average coalescence time for NEF outliers in the GA cluster. On the other hand, genes with old coalescence times in GA also displayed old coalescence times in NEF (Figure 7). This suggests that genes under balancing selection in GA were already under balancing selection before the split with Florida.

**Figure 7.**
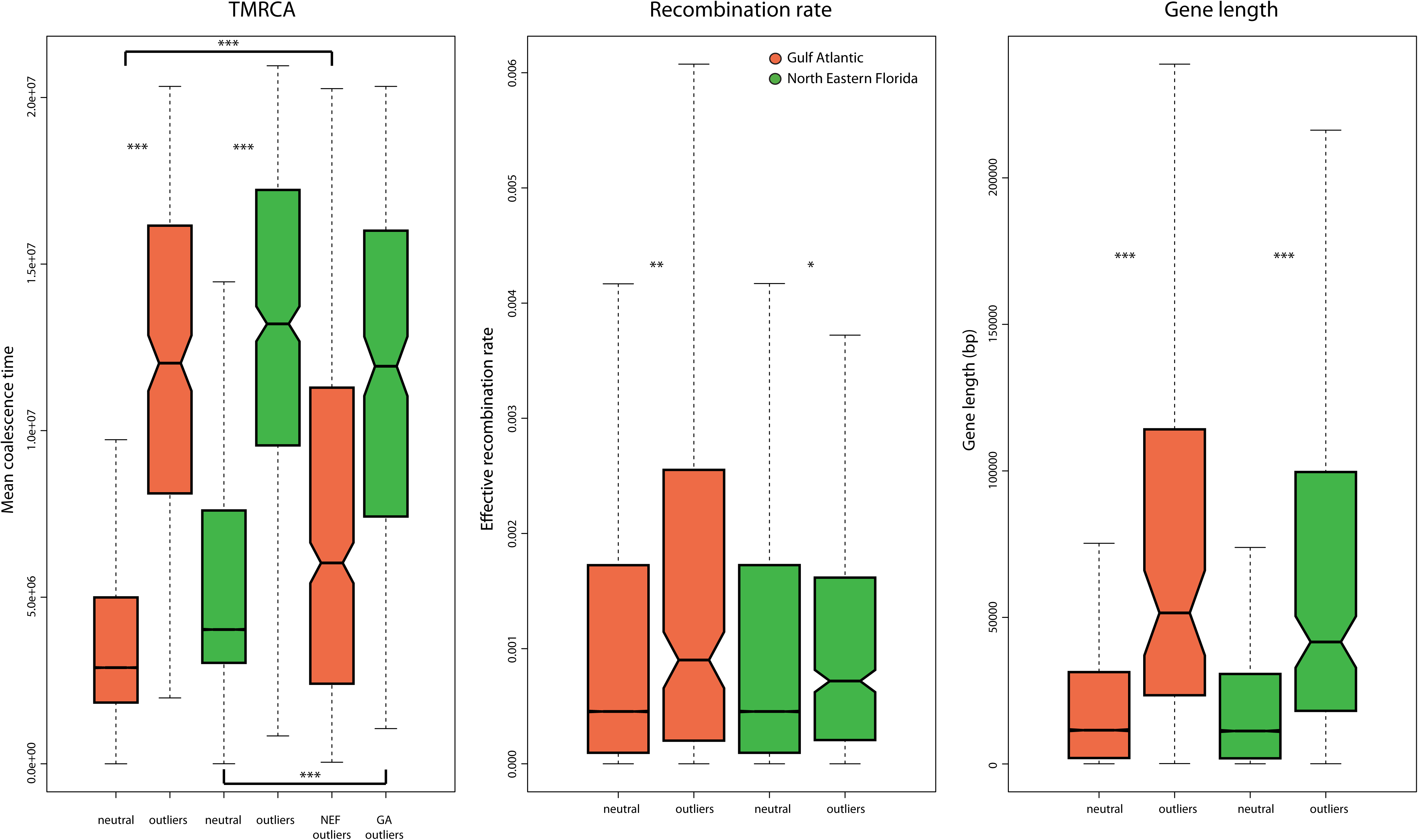
Properties of windows under balancing selection. For TMRCA, the two boxplots on the left correspond to the distributions of coalescence times in GA or NEF for windows classified as selected in NEF and GA respectively. ***: p-value<0.001, Wilcoxon tests. Only genes overlapping SNPs with a β score were included in the background set.

### Recombination and gene length

There was a clear effect of recombination and gene length on the likelihood for a gene to be classified as selected. Windows classified as under positive selection had significantly lower recombination rates when compared to genomic background (Figure 6). The average length of genes overlapping these windows was also longer compared to the set of non-candidate genes. Outlier windows for balancing selection displayed higher effective recombination rates on average, and genes in these windows were longer than in the genomic background (Figure 7).

We tested for each scan of positive selection whether genes belonging to GO categories of interest (Immune response, GO:0006955; somitogenesis and limb development, GO:0001756 and GO:0060173; Reproduction, GO:0000003; Behavior, GO:0007610) also displayed stronger evidence for selection while controlling for gene length and clustering (Table 3). We also investigated whether genes involved in response to cold displayed more signatures of selection than expected by chance, using a list of 1,375 genes highlighted in a recent transcriptomic study (Campbell-Staton et al. 2018). Genes belonging to the GO term “behavior” (GO:0007610) overlapped with windows classified as selected across the three tests for positive selection more often than expected by chance. These genes and “cold response” genes also displayed significantly higher eBPis and LSD scores. Behavioral genes did not display significantly lower recombination rates when compared to permuted genomic windows of the same size and clustered in the same way. This was not the case however for “cold resistance” genes, that displayed a significantly lower average recombination rate than expected by chance. However, the shift in statistics remained even after removing genes with low average recombination rates (i.e. falling in the bottom 25%).

## Discussion

We investigated signatures of positive and balancing selection in the green anole genome, focusing on populations that recently colonized temperate North America from Florida. This choice was made to take advantage of the strong differences in ecology and morphology between northern and Floridian populations (Goodman et al. 2013; Campbell-Staton et al. 2016), and to determine the genetic bases of adaptation to a drastically different environment. We used several state-of-the-art methods to detect selection in genomes, both to increase confidence in the set of candidate genes for selection but also to gain insight on the behavior of these methods and compare their outputs. While detecting significant overlap between different methods is a common challenge in GWSS (Pavlidis et al. 2012), we generally observed a good agreement. Strikingly, we detected signatures of positive selection at genes involved in adaptation to cold, but also at genes involved in behavior and neural development. These findings were robust to gene length, clustering and recombination.

### Colonization of novel environments requires metabolic and behavioral adaptation

Anoles are known to be good colonizers, and constitute a model to understand biological invasions (Kolbe et al. 2017; Tamate et al. 2017). In the green anole, there is strong evidence for a significant broadening of the occupied ecological niche compared to its Cuban sister species (Campbell-Staton et al. 2016). This broadening is associated with significant physiological changes in lizards living in a more temperate climate, such as larger body size (Goodman et al. 2013) and adaptation to cold (Campbell-Staton et al. 2016). We do observe signatures of selection that are consistent with these adaptations, with an enrichment for genes involved in metabolic process across most tests for positive selection (Sup. Tables, Table 1). For example, we find signatures of selection at *ITCH*, a gene involved in inflammation (Perry et al. 1998; Giamboi-Miraglia et al. 2015), homeostasis and hematopoiesis (Chen et al. 2001). Other genes are involved in coagulation, inflammation or plasminogen activation, such as *SERPINI2* (Xiao et al. 1999; Heit et al. 2013) or *TTC7* (Nüesch et al. 2018).

Among these genes also lie growth factors *(LTBP1, PDGFB, HBEGF)*, receptors to growth factors (e.g. *ACVR1)* or transcription factors (e.g. *RUNX2)* that may have been important to morphological and physiological adaptation. For example, *LTBP1* has been shown to control the development of limb buds in chicken (Lorda-Diez et al. 2010), and *RUNX2* displays important functions in both skeleton development (McGee-Lawrence et al. 2014) and circadian behavior (Reale et al. 2013). To better connect our results with functional studies performed on green anoles, we examined the distribution of statistics for selection at genes involved in cold tolerance (Campbell-Staton et al. 2018) and found these to be skewed towards higher values than expected by chance. This suggests that polygenic adaptation to colder environmental conditions played an important role during the initial stages of mainland colonization.

A remarkable aspect of our study is the finding that many genes related to nervous system development and behavior displayed signatures of positive selection in the branch leading to northern populations, a pattern that did not seem to be due to systematic biases in gene length, clustering or recombination rate. Anoles are an important model for behavioral studies (Lovern et al. 2004), and behavioral traits play a major role in explaining survival and adaptation to novel environmental conditions in these lizards (Losos et al. 2004; Lapiedra et al. 2018). In addition, terrestrial ectotherms display diverse behavioral strategies to regulate their body temperature, adapting to ambient environmental conditions that can fluctuate in time and space (Blouin-Demers et al. 2003; Bouazza et al. 2016). For example, green anoles from northern Tennessee remain active during cold winter months by aggregating in crevices where they find protection against cold temperatures (Bishop and Echternacht 2004).

We found signatures of positive selection at three well-known genes involved in exploratory behavior (GO:0035640), *DRD4, DEAF1* and *LSAMP*. *LSAMP* is a glycoprotein that controls the growth of limbic neurons and guides the projection of limbic fibers. In humans, the limbic system is a crucial component of the regulation of emotional behavior, learning and memory (Horton and Levitt 1988). *LSAMP*-KO mice display a high reactivity to novel stimuli and environments (Catania et al. 2008), and show an increase in the amount of risk-assessment. DRD4-KO mice are less prone to explore new environments (Dulawa et al. 1999), while *DEAF1-*KO mice exhibit stronger anxiety (Vulto-Van Silfhout et al. 2014). Evidence for a role of *DRD4* in behavior and dispersal is also found in several bird species (Fidler et al. 2007; Garamszegi et al. 2014), and is linked to adaptation to changing climate in a migratory songbird (Bay et al. 2018). In humans, this gene has been the focus of several behavioral studies, showing that polymorphism at this gene predicted individual differences in novelty-seeking (Ebstein et al. 1996) and impulsivity (Carpenter et al. 2011). These findings echo those from a previous report (Bourgeois et al. 2018a) that detected possible signatures of strongly male-biased dispersal in northern populations of green anoles. In anoles, male dispersal if often related to competition, showing increased dispersal when density increases and competition for females is stronger (Johansson et al. 2008). In *Anolis sagrei*, smaller males tend to disperse more and avoid competition from larger males (Calsbeek 2009). Overall, these results suggest that the recent northward colonization from Florida may have been facilitated by disruptive changes in behavior and increased exploration of unknown environments, possibly due to competition for resources.

### Balancing selection maintains diversity at immune and developmental genes

We detected signatures of long-term balancing selection at genes involved in immune response in mainland populations, a pattern commonly observed in vertebrates (Piertney and Oliver 2006). In general, coalescence times for regions under balancing selection in Florida were reduced in the northern populations. The fact that genes involved in immunity escaped this trend suggests an important role of biotic constraints in maintaining diversity even in bottlenecked populations. This confirms the resilience of this type of adaptive variation to demographic perturbations (Oliver and Piertney 2012; Bourgeois et al. 2018b).

We also found evidence of strong balancing selection at genes involved in appendage development and locomotion, nervous system development and learning (Table 2). For example, *RARB* is involved in the mesolimbic dopamine signaling pathway and regulates locomotory behavior (Krȩzel et al. 1998), and *ALX4* plays a critical role in the antero-posterior patterning of limbs (Kuijper et al. 2005). These signals were particularly strong in the NEF cluster. This finding is intriguing, and the exact nature of the selective pressures that may cause it remains elusive. However, a classical example of balancing selection maintaining morphological and behavioral polymorphism is found in the rock-scissor-paper dynamics of side-blotched lizards (Sinervo et al. 2001). A recent molecular study in bank voles has shown that sexual antagonism and density-dependence can lead to balancing selection at behavior genes (Lonn et al. 2017).

There is evidence for polymorphism in behavior and size in green anoles (Lailvaux et al. 2004), although the phenotypes may be age-dependent. Other behavioral traits such as headbob display are also polymorphic and change in frequency across populations depending on habitat and populations density (Bloch and Irschick 2006). It is worth noting that dominant young males tend to display higher jumping velocity and similar bite force than losing young males, but that this pattern is reversed in old males (Lailvaux et al. 2004). Such a trade-off mechanism where males having the highest reproductive success at a young age end up losing at an older age could maintain polymorphism at developmental and behavior genes, and deserves further investigation.

### Gene length and recombination impact the likelihood to detect genes under selection

Candidate windows for positive selection were mostly found in regions of lower recombination (measured as 4N*r in the NEF population) than in the rest of the genome. The length of the genes overlapping these windows was also longer than the length of non-selected genes. An interesting aspect is that genes involved in cold adaptation in anoles (Campbell-Staton et al. 2018) have a significantly lower average effective recombination rate than expected (Table 3). There are several interpretations to these trends. The first one is simply that positive selection is easier to detect in regions of low recombination because of stronger hitchhiking generating long, differentiated haplotypes. In addition, longer genes will overlap more windows, rising the odds to detect outliers. Moreover, false-positives are also expected to cluster in regions of lower recombination because of the expected long-distance correlation in allele frequencies. Another explanation lies in the theoretical expectation that genes under positive selection are also more likely to be found in regions of low recombination. Low recombination increases linkage between adaptive variants, a process that might explain negative correlations between differentiation metrics and recombination, although empirical evidence remains limited (Samuk et al. 2017). Further finer-grained studies focusing on multiple populations with varying degrees of gene flow and selective pressures may help distinguishing between these two explanations.

Genes under balancing selection were also longer than neutral genes, and their effective recombination rates were higher. This last feature is more likely to be explained by larger effective population sizes (4N) in NEF at these loci than by higher recombination rates (r). These locally larger effective population sizes are expected in the case of balancing selection (Charlesworth 2006).

### The need for genomic studies in a model for physiology and behavior

Genomic studies in anoles are still scarce, and only a couple of studies so far have used genomic data to answer questions about adaptation in this clade (Campbell-Staton et al. 2016; Tamate et al. 2017). Our results show that these approaches may have the potential to draw a more comprehensive picture of the links between genotype, phenotype and environment in vertebrates. We want to insist at this point that our tests were mostly aimed at targeting signatures of positive selection shared between northern clades. This means that several of our candidates will have been selected prior or concomitant to the split between northern clades and their expansion. Our set of candidates is therefore likely enriched in genes that may have been involved in the fast colonization of northern and novel environments, but the actual selective pressures at the time of selection remain difficult to assess. However, it seems likely that the ancestral population of northern clades was already found at the margin of Floridian environmental niche, where it could already develop adaptation to colder environments.

Our study is therefore set at a broader spatio-temporal scale than the study led by Campbell-Staton et al. (Campbell-Staton et al. 2016) which used explicit association tests with environmental features and may have had more power to detect genes specifically linked to adaptation to cold, especially in farther north populations. Further effort should assess how the reduction in genetic variation due to both positive selection prior colonization and bottlenecks impacted further adaptation to cold environment. Such an approach has the potential to shed light on the genetic bases of adaptation at coarse-grained and fine-grained spatio-temporal scales (Manel et al. 2010; Perrier and Charmantier 2018).

Although powerful, GWSS are not exempt of pitfalls (Li et al. 2012; Pavlidis et al. 2012). For example, demography and intrinsic properties of genomes such as variation in recombination rates have sometimes been overlooked in GWSS (Pavlidis et al. 2012; Haasl and Payseur 2016). Avoiding these issues requires working with species for which preliminary knowledge of the non-adaptive factors shaping genome variation is already available. In addition, caution should be exercised against overinterpretation and storytelling, as “genes that make biological sense” may be found even when randomly sampling the genome for candidates (Pavlidis et al. 2012). Our current and previous work has aimed at alleviating the issues stated above, by providing an account of the non-adaptive factors shaping genome variation (Bourgeois et al. 2018a), by comparing signatures of selection between candidate genes and genomic background, and by taking advantage of the knowledge of selective agents to interpret results from GWSS (Haasl and 2016).

We note that there are redundant patterns between GWSS in green anoles, such as the repeated finding that candidate genes for positive selection are involved in metabolism, muscular development and physiology (Campbell-Staton et al. 2016; Tamate et al. 2017; Tollis et al. 2018; Prates et al. 2018), even in extremely recent adaptation (Tamate et al. 2017). Although this is encouraging, we urge to apply caution since most developmental genes tend to be longer than other annotated genes in the green anole (Sup. Figure 1), which can make them easier to detect by GWSS. The temptation of favoring storytelling over cautious interpretation of the significance of such candidates can be strong given the importance of anoles in studies on physiology, morphology and behavior. To avoid this pitfall, future work should attempt to bring together functional and genomic frameworks to investigate the actual effect of variation at candidate genes for selection (Donihue and Lambert 2015; Haasl and Payseur 2016; Wadgymar et al. 2017).

Predictability in evolution is sometimes elusive, and quantifying the response of organisms to selection remains challenging (Pujol et al. 2018). In this context, molecular ecology and population genomics provide powerful tools to bring together phenotypic and genotypic information into an integrated framework. We believe that our study may contribute to a broader understanding of the evolutionary and functional mechanisms underlying the outstanding phenotypic diversity in anoles, and more generally vertebrates.

## Methods

### Sampling and Whole Genome Sequencing

Whole genome sequencing libraries were generated from 27 *Anolis carolinensis* liver tissue samples collected between 2009 and 2011 (Tollis et al. 2012), and *A. porcatus* and *A. allisoni* tissue samples generously provided by Breda Zimkus at Harvard University. A detailed summary of laboratory and bioinformatics procedures are already presented in (Bourgeois et al. 2018a). Sequencing depth was comprised between 7.22X and 16.74X, with an average depth of 11.45X.

The analysis included 74,920,333 variants with less than 40% missing data. Sequencing data from this study have been submitted to the Sequencing Read Archive (https://www.ncbi.nlm.nih.gov/sra) under the BioProject designation PRJNA376071. We extracted bioclimatic data for each locality from the WORLDCLIM database ((Hijmans et al. 2005) http://www.worldclim.org/version1), using the R packages raster and rgdal (Hijmans 2014; Bivand et al. 2015). PCA on this set of variables was performed in R.

### Genome scan using diploS/HIC

We used a recently developed machine-learning algorithm diploS/HIC (Kern & Schrider, 2018) to classify genomic windows as selected or not in northern clades. We followed a procedure similar in spirit to a previous study (Bourgeois et al. 2018b). We trained the algorithm using a set of 3,000 coalescent discoal simulations (Kern and Schrider 2016). We simulated 330kb windows divided into 11 subwindows. Hard and soft sweep examples consisted in windows with a sweep occurring in central 30kb subwindow. We sampled selection coefficients from a uniform prior such as 2Ns ~ (30,3000) for soft sweeps and 2Ns ~ (30,300) for hard sweeps, due to the stronger effect of hard sweeps on diversity. For soft sweeps, we used uniform priors on the initial frequency of the adaptive variant of (0,0.2). We assumed a mutation rate of 2.1×10^−10^ per site per generation, a generation time of one year (Tollis and Boissinot 2014) and a recombination rate of 8.25×10^−11^/site/generation estimated in the NEF cluster from a previous study (Bourgeois et al. 2018a). We used priors for present effective population sizes between twice lower and five times higher the values estimated from a SMC++ analysis (Figure 1, (Terhorst et al. 2016; Bourgeois et al. 2018a)) to consider the uncertainties in current size estimates. We used a truncated exponential prior for recombination rates with an average of 8.25×10^−11^ and a maximum value of 1.8×10^−9^. We conditioned on sweep completion occurring between 500,000 generations ago and shortly before northern clusters diverged from each other, around 150,000 generations ago. We allowed for variable gene flow between northern populations after their split (0<4N_0_m<0.5 with N_0_=1,475,000, the current effective population size of the GA cluster). Categorization of 30kb windows as sweeps, linked or neutral was then performed on the actual dataset, removing scaffolds shorter than 330 kb and sex-linked regions. diploS/HIC assigns a probability to each window for being a sweep, linked to a sweep, or neutral. We took advantage of this probability of assignation to improve the false positive rate by retaining candidate windows classified as sweeps only if they displayed a probability < 10% of being neutral.

### Genome scan using LSD

Individuals were first phased using BEAGLE v4.0 with default options (Browning and Browning 2007). Haplotypes trees were computed over non-overlapping 1,250 SNPs windows with PhyML v3.1 (Guindon et al. 2010) using the script phyml_sliding_windows.py available at https://github.com/simonhmartin/genomicsgeneral. We chose to use 1,250 SNPs windows instead of windows of fixed lengths to get an equivalent amount of information across windows and increase resolution in regions of high recombination where SNP density and diversity is higher (Bourgeois et al. 2018a). The LSD algorithm was run over the 5 genetic clusters identified in a previous study (Figure 1, (Bourgeois et al. 2018a)), using northern clades as the focal branch in which to detect selection (noted A1 and A2, see https://bitbucket.org/plibrado/LSD) while the Floridian clade NEF was identified as the sister branch to northern clades. The trees were rerooted by the algorithm using *Anolis allisoni* as an outgroup. We considered the windows ranking in the top 1000 LSD scores as candidates for positive selection and extracted overlapping genes (Ensembl release 90, genome version: AnoCar2.0) using the intersect function in bedtools v2.25.0 (Quinlan and Hall 2010).

### Genome scan using BAYPASS

Overall divergence at each locus was first characterized using the *X^T^X* statistics, which is a measure of adaptive differentiation corrected for population structure and demography. BAYPASS offers the option to estimate an empirical Bayesian p-value (eBPis) which can be seen as the support for a non-random association between alleles and specific populations. We computed eBPis over the top 5% *X^T^X* outliers to determine their level of association with northern populations. BAYPASS was run using default parameters under the core model. We considered a SNP as a candidate for selection in northern populations when belonging to the top 1% eBPis. We divided the genome in 5kb windows and retained those with at least three outlier SNPs as candidates for positive selection. Genes overlapping these windows were extracted using bedtools v2.25.0.

### Genome scan for long-term balancing selection

We used the software Betascan to estimate the β score, a statistic that detect clusters of alleles segregating at similar frequencies across windows of a size that is predefined by the user (Siewert and Voight 2017). To determine the optimal window size in which signals of balancing selection should be detectable, we estimated the expected length distribution of selected haplotypes using the formula provided in (Siewert and Voight 2017). This distribution is exponential with rate parameter T*ρ with T the time since balancing selection and ρ the recombination rate. Since our goal is to detect long-term balancing selection in the green anole, we aim at still being able to detect events having occurred around the split with its sister species, *Anolis porcatus*, from which it diverged between 6 and 12 million years ago (Glor et al. 2005). Using T=12 million generations and ρ =8.25×10^−11^, the 95^th^ quantile on either side of a selected SNP is 3026 bp. We therefore decided for the sake of simplicity to use a window size of 2.5 kb on each side of each focal SNP (for a total size of 5kb, option -w 5000). We did not compute β scores for SNPs with a folded allele frequency below 0.1 (−m 0.1) and rose the “-p” parameter to 50 instead of the default 20 to ensure strong correlations between allele frequencies for the SNPs surrounding candidate loci. We focused our analyses on Gulf Atlantic and Central Florida to minimize confounding effects of population structure and to assess the fate of genes under balancing selection following bottleneck and adaptation. We divided the genome in 5kb windows and retained those with at least three outlier SNPs (i.e. in the top 0.1% β score) as candidates for positive selection.

### Ancestral recombination graphs

Genes under positive selection should display reduced coalescence times in the northern clades but should not display such reduction at the species scale. On the other hand, genes under balancing selection should display older haplotypes and therefore older coalescent times than the genomic background. We used the software ARGWeaver to confirm that our tests of selection displayed results consistent with these expectations and provide a quantification of the timing of events of selection (Rasmussen et al. 2014). We included in the analysis a subset of 12 individuals with high sequencing depth and covering the species range (between 2 and 3 individuals from each cluster, Sup. Table 8) to limit computation time. Individuals were first phased using BEAGLE v4.0 with default options but did not infer missing data (Browning and Browning 2007). We took a mutation rate of 2.1×10^−10^/bp/generation and a recombination rate of 8.25×10^−11^. We used a prior effective population size of about 1,000,000 which is of the same order of magnitude than previous estimates for all green anole populations obtained from targeted sequencing and whole genome data (Tollis and Boissinot 2014; Manthey et al. 2016; Bourgeois et al. 2018a). ARGWeaver reconstructs ancestral recombination graphs and therefore accommodates variable recombination rates, effective population sizes and genealogies along the genome. The algorithm was run for 1000 iterations, using 15 discretized time steps and a maximum coalescence time of 50 million generations. We then extracted times since the most recent common ancestor (TMRCA) for each non-recombining block in GA, CA and NEF clusters, as well as for the whole set of 12 individuals. To compare coalescence times between windows classified as selected or neutral (e.g. LSD windows or 5kb windows), we averaged values using the package regioneR (Gel et al. 2015), weighing by the length of non-recombining blocks.

### GO-enrichment

We tested for enrichment of overrepresented Gene Ontology (GO) terms associated with the genes found in outlier regions using the PANTHER analysis tool available on the Gene Ontology Consortium website (Fisher tests, www.geneontology.org, last accessed October 2018; (Blake et al. 2015)). We took care of adjusting the gene universe depending on the test for selection by contrasting the set of candidate genes to the set of genes actually covered by each test. We tested for enrichment of any biological function. We did not apply correction for multiple testing for several reasons: i) since this approach is exploratory, we wanted to detect any interesting trend in the dataset that could then be further explored; ii) we focused on the clearest signatures of positive selection, limiting the number of outlier genes to a few hundred across several potentially relevant GO terms. Instead we present the first 50 most significant GO terms for each test and isolate candidate genes falling in GO categories of interest. Lists of all genes identified by each approach are available as Supplementary material (Sup. Tables 9 to 14).

### Permutation tests and correction for clustering, gene length and recombination

We used bedtools intersect and the package regioneR to estimate average lengths and recombination rates for genes found in outlier regions and compare them to genomic background. We also used regioneR to assess whether the distribution of statistics for positive selection (LSD and eBPis) was significantly shifted in GO categories that are relevant in anoles. We used both z-scores and permutation p-values to estimate the significance of this shift. More specifically, we used the circularRandomizeRegions function, that takes a set of genomic intervals (here candidate genes from a given GO term) and applies a random spin to each chromosome. This approach therefore maintains clustering and gene length in the pseudo-observed dataset. We used 1000 permutated datasets as a random expectation for the average values of recombination rates, LSD and eBPis scores, as well as for the number of overlaps between genes of interest and windows classified as selected by all three approaches.

## Supporting information

## Acknowledgements

We are grateful to Breda Zimkus from the Museum of Comparative Zoology Cryogenic Collection in Harvard and J. Rosado from the Herpetology Collection for providing the samples of *Anolis porcatus* and *Anolis allisoni.* We thank Justin Wilcox, Jacobo Reyes-Velasco, Sandra Goutte and Sebastian Kirchhof for their comments on the manuscript. We thank Marc Arnoux from the Genome Core Facility at NYUAD for assistance with genome sequencing. This research was carried out on the High Performance Computing resources at New York University Abu Dhabi. This work was supported by New York University Abu Dhabi (NYUAD) research funds AD180 (to S.B.). The NYUAD Sequencing Core is supported by NYUAD Research Institute grant G1205-1205A to the NYUAD Center for Genomics and Systems Biology.

---

**Supplementary Figure 1:** Distribution of lengths for genes belonging to several GO categories of interest in green anoles. Only genes with a GO annotation for biological function were included (N=18,523). Legend: only GO:0008152, genes assigned to metabolic process (but not anatomical structure development, N=5307); only GO:0048856, genes assigned to anatomical structure development (but not metabolic process, N=1464); GO:0008152+GO:0048856” genes assigned to both metabolic process and anatomical structure development (N=1131). All p-values< 2.2×10^−16^ (Wilcoxon tests comparing genes of interest with the background).

**Supplementary Table 1:** Confusion matrix for diploS/HIC. Percentages of assignation to each category (in columns) are indicated. Each row corresponds to a set of 3,000 discoal simulations.

**Supplementary Table 2:** GO enrichment analysis for genes classified as selected by diploS/HIC.

**Supplementary Table 3:** GO enrichment analysis for outlier genes in LSD analysis

**Supplementary Table 4:** GO enrichment analysis for outlier genes in BAYPASS analysis.

**Supplementary Table 5:** GO enrichment analysis for genes overlapping candidate windows for positive selection across three methods.

**Supplementary Table 6:** GO enrichment analysis for outlier genes in Betascan analysis (GA).

**Supplementary Table 7:** GO enrichment analysis for outlier genes in Betascan analysis (NEF).

**Supplementary Table 8:** Individuals included in the ARGWeaver analysis and their coordinates.

**Supplementary Tables 9-14:** List of outlier genes for LSD, diploS/HIC, BAYPASS, these three methods, the Betascan analysis in GA and the Betascan analysis in NEF.

