## Supplementary figures and images for "Selection at behavioral, developmental and metabolic genes is associated with the northward expansion of a successful tropical colonizer"

### Supplementary file 1

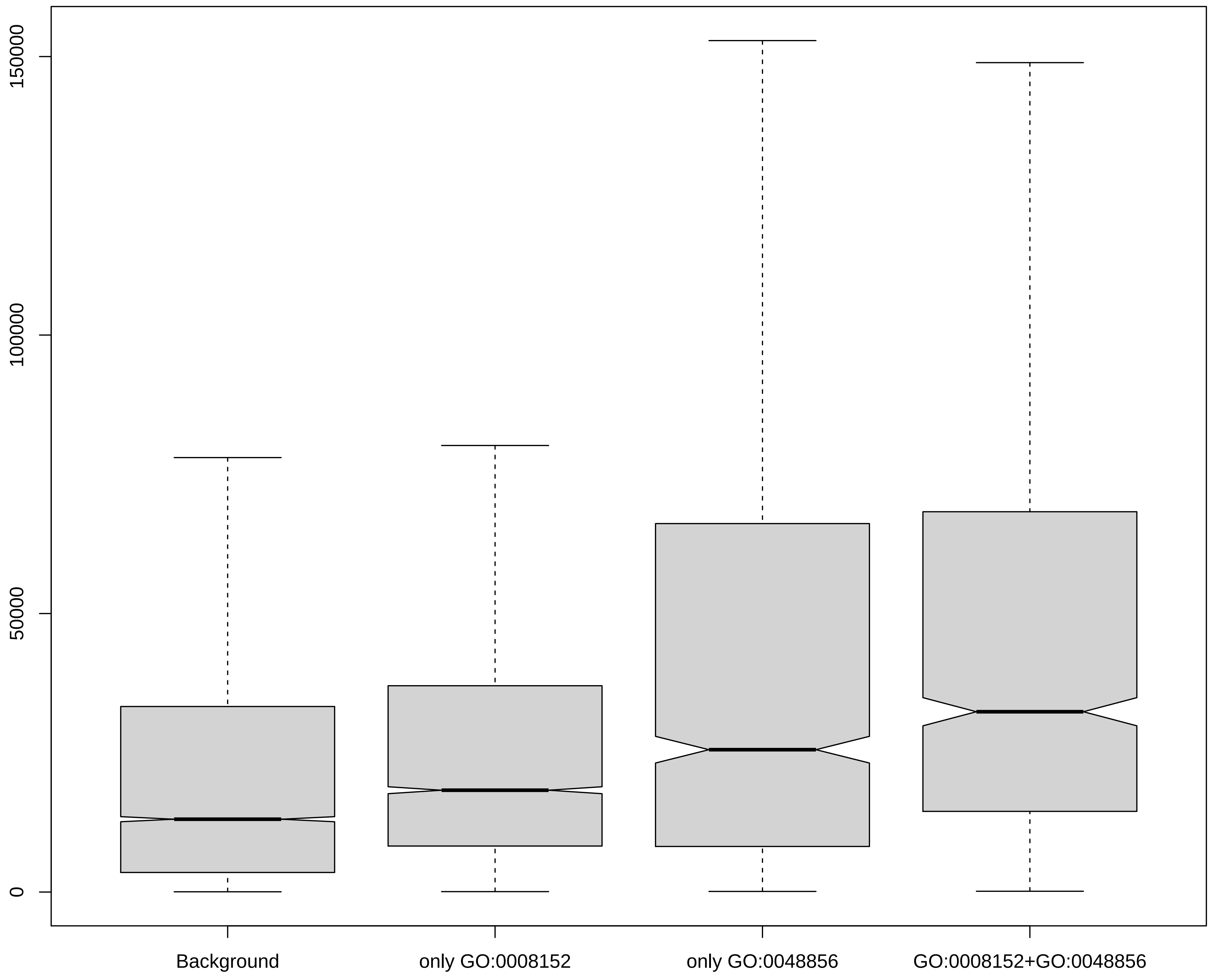
